# Remotely Presenting Alcohol-predicting Cues Avoids Confound of Experimenter as First Cue and Reveals Sex-specific Behaviors that Predict the Rate and Amount of Alcohol Consumption

**DOI:** 10.64898/2026.08.07.743581

**Authors:** Seth A. David, Dominic A. Furlano, Monserrat Orozco, David N. Linsenbardt

## Abstract

Understanding the neurobiological systems that regulate alcohol cue-induced craving is of utmost importance for the development of novel intervention strategies for alcohol use disorders (AUDs). However, although a human experimenter is required to conduct alcohol self-administration studies in the lab, the cues associated with the experimenter are seldom if ever factored into the experimental design. Thus, although we have learned much to date about alcohol cue-induced behavior and neurobiology, and in particular about discrete cues presented many times throughout a single daily alcohol self-administration session, we know relatively little about how responses to alcohol availability cues might predict subsequent alcohol consumption. For the current experiment, mice were exposed daily to auditory cues that preceded 2 hours of alcohol or water access using drinking-in-the-dark (DID) methods. An additional control group experienced cues but were not otherwise manipulated. Importantly, cues were initiated remotely from outside the animal facility, avoiding the experimenter being the first cue predicting ethanol availability. Head direction, location in the home cage, and movement velocity were the primary variables on interest. Surprisingly, during the cue period, there were no significant differences between groups in any of these measures, despite meaningful alterations over days. However, we observed many significant correlations between behaviors and drinking variables. First, we observed significant positive associations between ambulatory velocity during cues and subsequent total alcohol (R^2^=0.14; p<0.0001) and total water (R^2^=0.12; p=0.0002) consumption, but only in females. We also observed a significant positive relationship (R^2^=0.25; p<0.0001) between the amount of time oriented toward the sipper port during the auditory cues and the average *rate* of subsequent alcohol consumption (i.e. front-loading), but only in females. In males, head direction was found to be positively associated with subsequent total water consumption (R^2^=-0.21; p<0.0001), but not alcohol (R^2^=-0.01; p=0.2267). We also observed a significant *negative* relationship (R^2^=-0.15; p<0.0001) between proximity to the sipper during the cue period and subsequent total 2-hour alcohol intake in males. Although these associations were modest in strength, they suggest potential sex-specific behavioral predictors of alcohol consumption that are regulated by different neural dynamics.

## 1. Introduction

Excessive alcohol consumption poses a major public health concern in the United States, contributing to approximately 178,000 deaths and an estimated $249 billion in economic costs annually (*CDC*, 2010). This toll is driven in large part by heavy and binge drinking patterns. In 2022, 6.9% of adults reported heavy drinking, defined as consuming more than 14 drinks per week for men and more than 7 for women. Additionally, 16.9% reported binge drinking, defined as consuming 5 or more drinks in a single occasion for men and 4 or more drinks in the same time frame for women (*CDC*, 2023).

Both chronic heavy drinkers and episodic binge drinkers are frequently exposed to recurring environmental and sensory stimuli during and prior to alcohol use. These stimuli, often complex, highly individual combinations of sights, sounds, locations, and odors, become conditioned cues in some individuals that elicit alcohol cravings (Witteman et al., 2015). For instance, cues previously paired with even low doses of ethanol have been shown to rapidly elicit both physiological arousal and subjective reports of craving (Field & Duka, 2002). Additionally, cue-induced cravings have been observed to emerge either before or during initial alcohol intake, depending on individual sensitivity to alcohols subjective effects (i.e. feeling buzzed; Kohen et al., 2023), underscoring the individualized nature of the role of cues in driving alcohol seeking.

Preclinical models have been instrumental in determining the influence of alcohol-associated cues on behavior. Prior studies have explored these cues in the context of relapse (Valyear et al., 2023; Cofresí et al., 2017), Pavlovian-instrumental transfer (Corbit & Janak, 2007; Delamater & Holland, 2008), and cue-induced approach behaviors such as orienting toward the alcohol receptacle (Lamb et al., 2020; Cofresí et al., 2018; Cofresí, Grote, et al., 2019; Cofresí, Monfils, et al., 2019). Additional work has established important distinctions between discrete and contextual cues (Valyear et al., 2020; Valyear et al., 2023; Sciascia et al., 2015) and between sign– and goal-tracking responses (Krank, 2003; Krank et al., 2008; Madayag et al., 2017; Villaruel & Chaudhri, 2016). However, there remains limited insight into how cues predicting alcohol availability may elicit behavioral responses capable of predicting subsequent alcohol consumption. To this end, Cofresi and colleagues (Cofresí et al., 2018; Cofresí, Monfils, et al., 2019) found that approach behavior toward an alcohol sipper predicted blood ethanol concentrations in a delay-conditioning task. Nonetheless, like most if not all self-administration studies, subjects were retrieved from their vivarium by experimenters immediately prior to alcohol availability, but were not observed during this time period for behavioral responses that might be indicative of craving and/or that predict subsequent alcohol consumption variables.

The pivotal role of environmental context in shaping learning and memory for alcohol-paired cues is well established (Chaudhri et al., 2010; Remedios et al., 2014; Valyear et al., 2017; Faria et al., 2008; Bianchi et al., 2024). Contextual cues influence both the development and expression of responses to discrete cues. For example, when conditioning occurred in a distinct environmental context, mice did not respond to the same cue in a different context (Remedios et al., 2014; Bianchi et al., 2024). This has important implications for interpreting cue-elicited behavior in preclinical settings, as it becomes difficult to disentangle the role of discrete versus contextual cues, particularly in experiments using chambers that do not explicitly investigate contextual conditioning. Although operant chambers provide precise experimental control, they create distinct environmental conditions that promote context-specific learning, and interaction with the experimenter serving as an initial cue, complicating the interpretation of discrete cue effects and limiting reproducibility. This issue extends to clinical applications, where extinction-based treatments are often conducted in settings that differ greatly from real-world cue exposure, potentially reducing their effectiveness.

To address one of the most difficult challenges in these types of studies, we modified a well-known binge-drinking model in which mice were exposed to auditory cues that preceded and accompanied ethanol or water access. Importantly, by using remote-controlled cue presentation, we minimized experimenter interaction, avoiding the experimenter being the first and potentially most salient cue predicting ethanol availability, and reducing the role of additional/novel contextual cues by experimenting entirely within the home cage. This exploratory study aimed to identify behavioral responses to alcohol-paired cues that might predict excessive alcohol consumption. Our approach emphasizes precise temporal alignment of behavior with cue exposure in a more scalable setting, which is particularly important as neuroscience increasingly relies on temporally precise tools like optogenetics and fiber photometry.

By characterizing cue-elicited behaviors in a low-interaction, home-cage setting, this work seeks to reduce contextual and experimenter-related confounds and enable more accurate modeling of motivational states. Ultimately, this study seeks to lay the groundwork for closed-loop systems in which specific behaviors predictive of high-risk alcohol consumption trigger real-time neural interventions (i.e. optogenetics), to reduce subsequent drinking. Such approaches represent a critical step toward mechanistic, temporally precise models of addiction and other maladaptive behavioral states.

## 2. Methods

### 2.1 Subjects & Design

In the present study, 48 C57BL/6 mice split in half by sex were maintained on a 12/12 reverse light-dark cycle and provided food (LabDiet, 5L0D – PicoLab® Laboratory Rodent Diet) ad libitum. After an acclimation period of 7 days, the mice were pseudo-randomly assigned to treatment groups based on their total distance travelled during the last two days of acclimation, controlling inherent ambulatory differences between groups. Following three additional days in new cage positions, mice underwent drinking-in-the-dark (DID; (Rhodes et al., 2005)) for 15 days, during which they received a single volumetric drinking monitoring sipper (VDM), providing temporally accurate drinking volumes, dispensing either water or 20% alcohol during hours 3-5 of the dark cycle. An additional “naïve” group served as a negative control and never received VDM sippers.

Mice were single housed in standard amber shoebox plexiglass cages with custom made clear lids, with eight ∼8mm holes on each corner and centered along each edge. A single fluid port was available in which standard water sippers were available for all fluid-assigned mice for 22 hours a day, and for naïve mice 24/7. Cages were arranged in blocks of four, with a single camera for each block. The location of the mice in the testing room was counterbalanced for sex and treatment.

### 2.2 Cue Presentation

Fifteen minutes prior to the first mouse receiving a VDM sipper, a discrete one-minute auditory cue was presented via the flushing of the VDM sippers, which produced 48 simultaneous and continuous loud clicking/pumping noises (∼150 total). To minimize experimenter presence, flushing was initiated using a remote computer interface. Following one-minute of initial sipper flushing with no experimenter presence, the experimenter then entered the room and tested the sippers for proper function, producing additional exposure to clicking, and lasting approximately an additional 14 minutes. This provided in total a 15-minute cue period preceding VDM availability.

### 2.3 Behavioral Measures/Statistics

In addition to obtaining temporally accurate drinking volumes, we continuously tracked the animals 24/7 in real time using ANYMAZE (Stoelting Co., St. Louis). This provided information on velocity, head direction, and distance from the sipper. Data were managed in python and analyzed using GraphPad Prism Version 10. Drinking data were first analyzed using 2-way ANOVAs, followed by linear regressions and one-sample t-tests for front-loading. Cue-period behaviors were first analyzed using 3-way ANOVAs, which were followed up by 2-way ANOVAs with Dunnett’s multiple comparisons tests. Subsequently, we ran regression analysis, paired t-tests, and correlations with drinking patterns

## 3. Results

### 3.1 Drinking

The use of volumetric drinking monitoring sippers (VDMs) enables the monitoring of drinking patterns with high temporal specificity. This was leveraged to consider total volume consumed in addition to front-loading, a behavior defined by the consumption of a disproportionately large volume immediately upon fluid presentation compared to throughout the whole drinking period (Ardinger et al., 2022). While there is debate on the specific window to define “immediately,” an accepted practice is to compare the first 15-minutes to the entire 2-hour drinking period (Linsenbardt & Boehm, 2014; Linsenbardt & Boehm, 2015; Wilcox et al., 2014; Ardinger et al., 2020). Representative examples of average drinking patterns from alcohol and water consuming animals on days 1 and 15 can be seen in Figure 1A-D.

**Figure 1.**
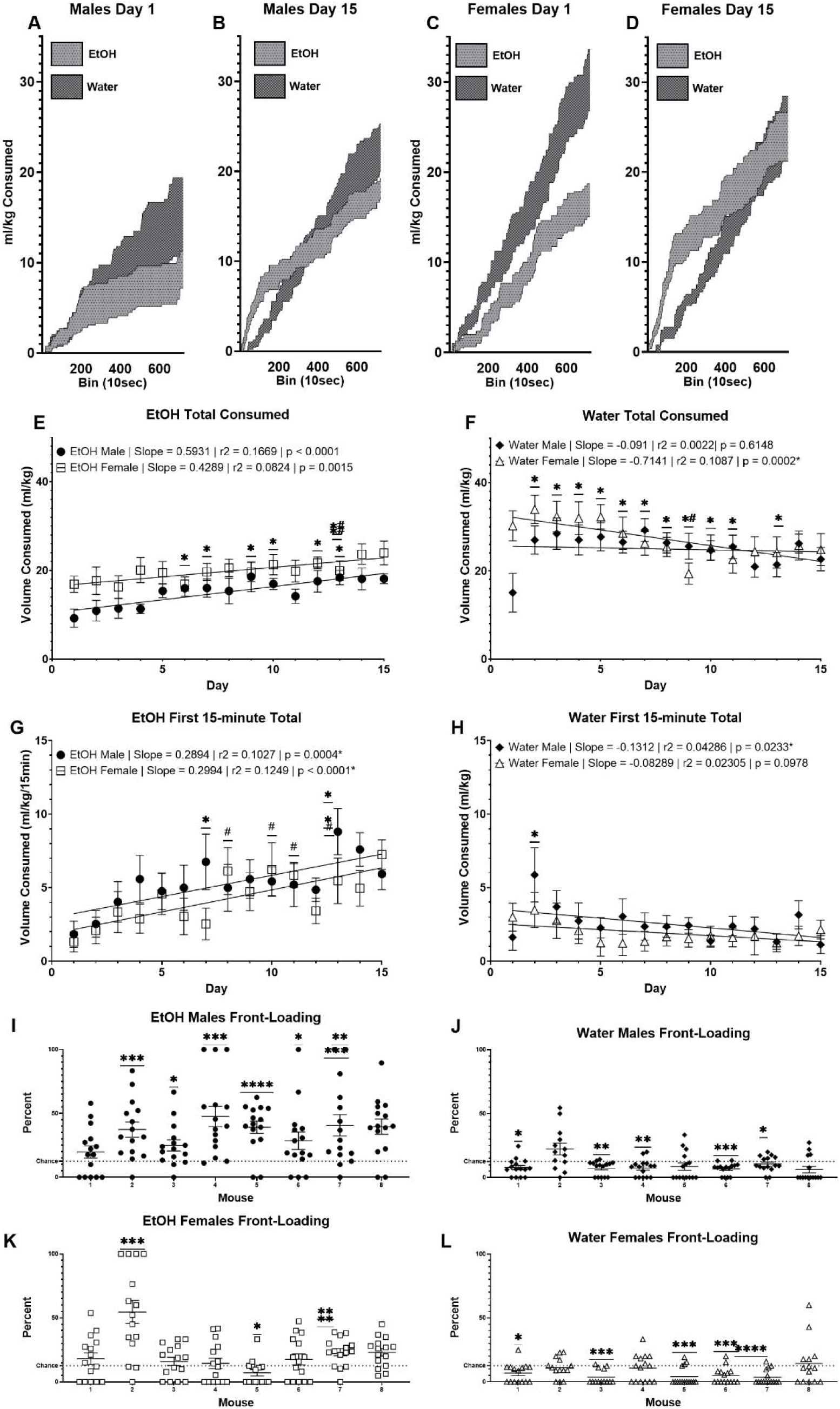
Fluid consumption profiles over the course of 15 consecutive days of DID. (A-D) representative images of volume consumed on days 1 and 15 throughout the 2-hour DID session. (E-F) Total volume consumed during DID on all days for ethanol-(E) and water-(F) assigned mice, split by sex. “*” represents males and “#” represents females in a 2-way ANOVA Dunnett’s multiple comparisons test comparing each day to day 1. (G-H) Volume consumed during the first 15-minutes of DID for ethanol (G) and water (H), split by sex. (I-L) One-sample t-tests of the percent of the total volume consumed during the first 15 minutes of DID, compared to chance levels of 12.5%.

Analysis of the total volume consumed for the entire 2-hour drinking period for the first 15 days was first analyzed using a three-way ANOVA. This revealed a significant main effect of treatment (DF = 1, F = 17.37, p = 0.0003), in addition to day x treatment (DF = 16, F = 5.806, p < 0.0001) and day x sex interactions (DF = 16, F = 2.477, p = 0.0013). Subsequent linear regressions elucidated the driving forces of these interactions, with alcohol-consuming mice of both sexes displaying significantly increases in consumption across days (Fig 1.E). Additionally, water females were the only group to significantly decrease consumption, while water males remained relatively stable (Fig 1.F). During the first 15-minutes of fluid access from VDM sippers, alcohol consuming mice of both sexes significantly increased consumption (Fig 1.G), while female water controls remained stable, and male water controls significantly decreased consumption (Fig 1.H).

If drinking rate were consistent across each of the 8, 15-minute drinking epochs that encompass the entire 2-hour session, 12.5% of total fluid would be consumed in each epoch. Thus, drinking more than 12.5% in the first 15 minute bin we defined as frontloading. T-test comparisons to this 12.5% threshold for each subject revealed that 7/8 Alcohol males frontloaded on average (Fig 1.I), whereas 3/8 Alcohol females (Fig 1.K) engaged in significant levels of front-loading across 15 days of drinking. By contrast, 5/8 water consuming mice of each sex consumed significantly *less* than chance during the first 15 minutes of the 2-hour drinking period (Fig. 1J,L).

### 3.2 Velocity

Due to large variability in individual subject velocity during initial baseline days, we used normalized velocity values, which were calculated as the percent change in ambulation during the 15-minute cue period preceding sipper availability compared to the mouse’s average velocity during the entire dark (active) period. We restricted our analysis to days 1-14 because experimenters remained present in the room following the DID session on day 15 which had the potential to confound results. Representative examples of average normalized velocity patterns from all animals on days 1 and 14 can be seen in Figure 2A-F.

**Figure 2.**
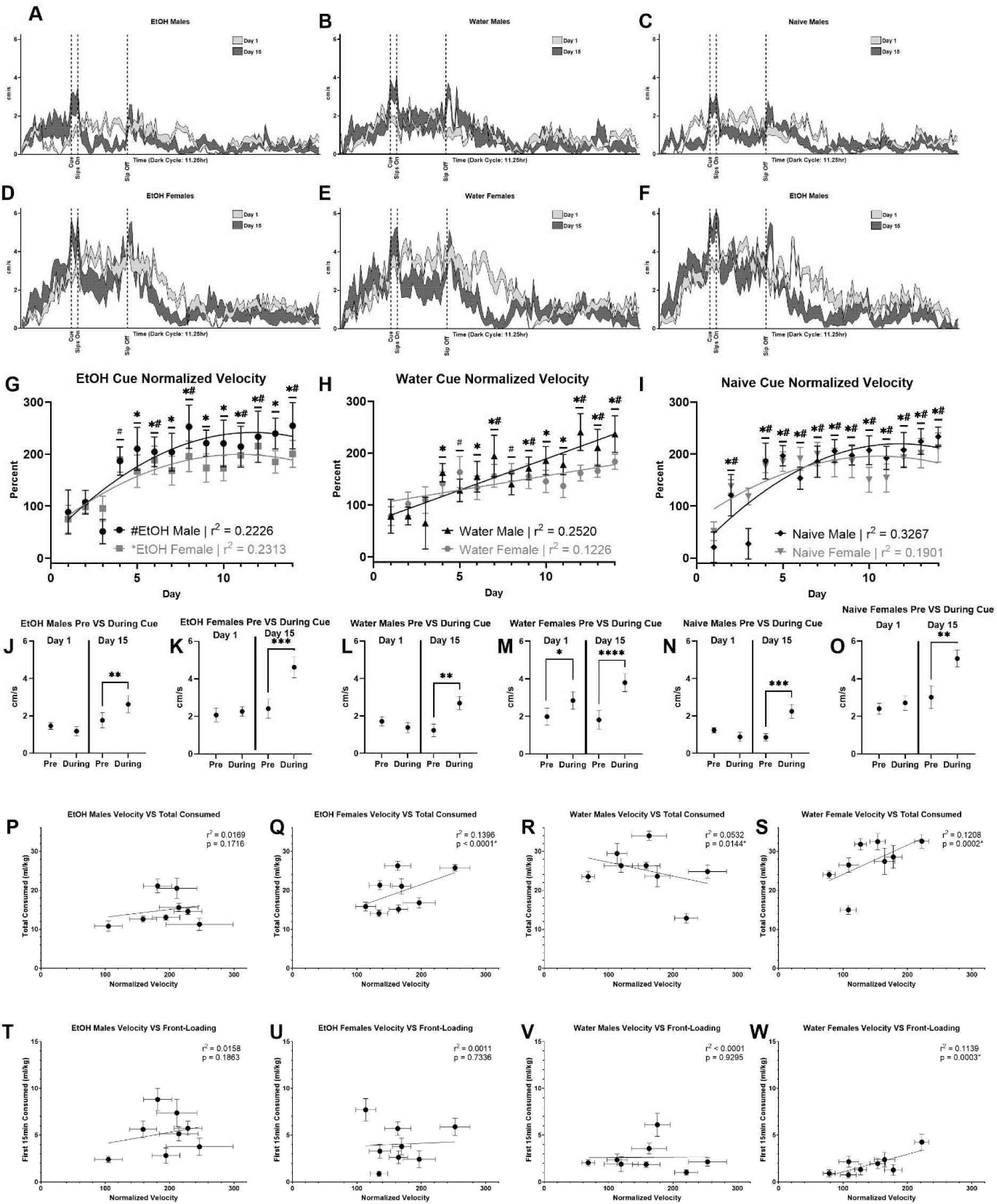
Impact of cues on ambulatory velocity and relationship to fluid consumption. (A-F) representative images of velocity throughout the dark (active) cycle on days 1 and 15, split by treatment and sex. (G-I) The percent increase in velocity during the 15-minute cue period relative to the average velocity during the entire dark cycle (((cue velocity – baseline velocity)/baseline velocity) *100). “*” represents males and “#” represents females in a 2-way ANOVA Dunnett’s multiple comparisons test comparing each day to day 1. (J-O) Paired t-tests of raw velocity during the 15-minutes prior to the cue and during the 15-minute cue on day 1 and day 15, split by treatment and sex. (J) EtOH Males Day 1: df = 7, t = 1.711, p = 0.1309; Day 15 df = 7, t = 3.718, p = 0.0075. (K) EtOH Females Day 1: df = 7, t = 0.5576, p = 0.5945; Day 15: df = 7, t = 6.378, p = 0.0004. (L) Water Males Day 1: df = 7, t = 1.38, p = 0.21; Day 15: df = 7, t = 4.125, p = 0.0044. (M) Water Females Day 1: df = 7, t = 3.284, p = 0.0134; Day 15: df = 7, t = 7.885, p < 0.0001. (N) Naïve Males Day 1: df = 7, t = 1.647, p = 0.1437; Day 15: df = 7, t = 7.056, p = 0.0002. (O) Naïve Females Day 1: df = 7, t = 1.565, p = 0.1616; Day 15: df = 7, t = 4.074, p = 0.0047. (P-W) Double-error-bar correlations between normalized velocity, total volume consumed, and first 15-min of DID volume consumed split by treatment and sex.

A three-way ANOVA revealed a significant main effect of day (DF = 13, F = 26.18, p < 0.0001) and a day x sex interaction (DF = 13, F = 2.334, p = 0.005). While there was no significant main effect of treatment, Dunnett’s multiple comparisons tests revealed that all groups significantly increased their velocity during the cue as the experiment progressed (Fig. 2. G-I). These treatment-independent increases indicate that the cues were salient enough to produce sensitization regardless of predictive value.

In addition to an increase in velocity during the cue relative to the entire lights off period, mice of all treatments and sexes also showed sensitization to the cue as compared to the 15-minutes immediately prior to its presentation on day 15 (Fig 2.J-O; p < 0.05). Only water consuming females showed a significant increase in velocity during the cue as compared to immediately prior to its presentation on day 1 (Fig 2.M; p = 0.0134).

Given the robust sensitization regardless of the fluid received, we were unsurprised to find that velocity was not uniquely predictive of the amount consumed in the first 15-minutes or the entire 2-hour drinking period when comparing alcohol and water mice. However, there were differences between sexes in the predictive validity of velocity on drinking patterns. In males, the only significant correlation was in total consumed within the water treatment (Fig 2.R; r2 = 0.0532, p = 0.0144), and was notably weak. In females, however, velocity was predictive of total volume consumed in both alcohol (Fig 2.Q; r2 = 0.1396, p < 0.0001) and water mice (Fig 2.S; r2 = 0.1208, p = 0.0002). Velocity was also significantly correlated with the first 15 minutes of water consumption in females (Fig 2.W; r2 = 0.1139, p = 0.0003). Altogether, these data suggest the degree of cue-induced increases in velocity provides some utility in predicting fluid consumption patterns.

### 3.3 Head Direction

The next behavior considered was head direction during the cue period. More specifically, the percentage of time spent oriented towards where the VDM sipper *will be* placed, which is where their normal non-VDM sipper lay throughout the other 22 hours of the day. A 3-way ANOVA revealed a main effect of treatment (DF = 2, F = 3.613, p = 0.0357); main effects of sex (DF = 1, F = 3.931, p = 0.054) and day (DF = 14, F = 1.704, p = 0.0508) were marginally significant. Linear regression revealed the only group to experience a significant change in time spent oriented toward the sipper during the cue across the first 15 days were the water males (Fig. 3A; slope = –0.2861, r2 = 0.0417, p = 0.0253); no changes in head direction over days were observed in females (Fig.3B).

**Figure 3.**
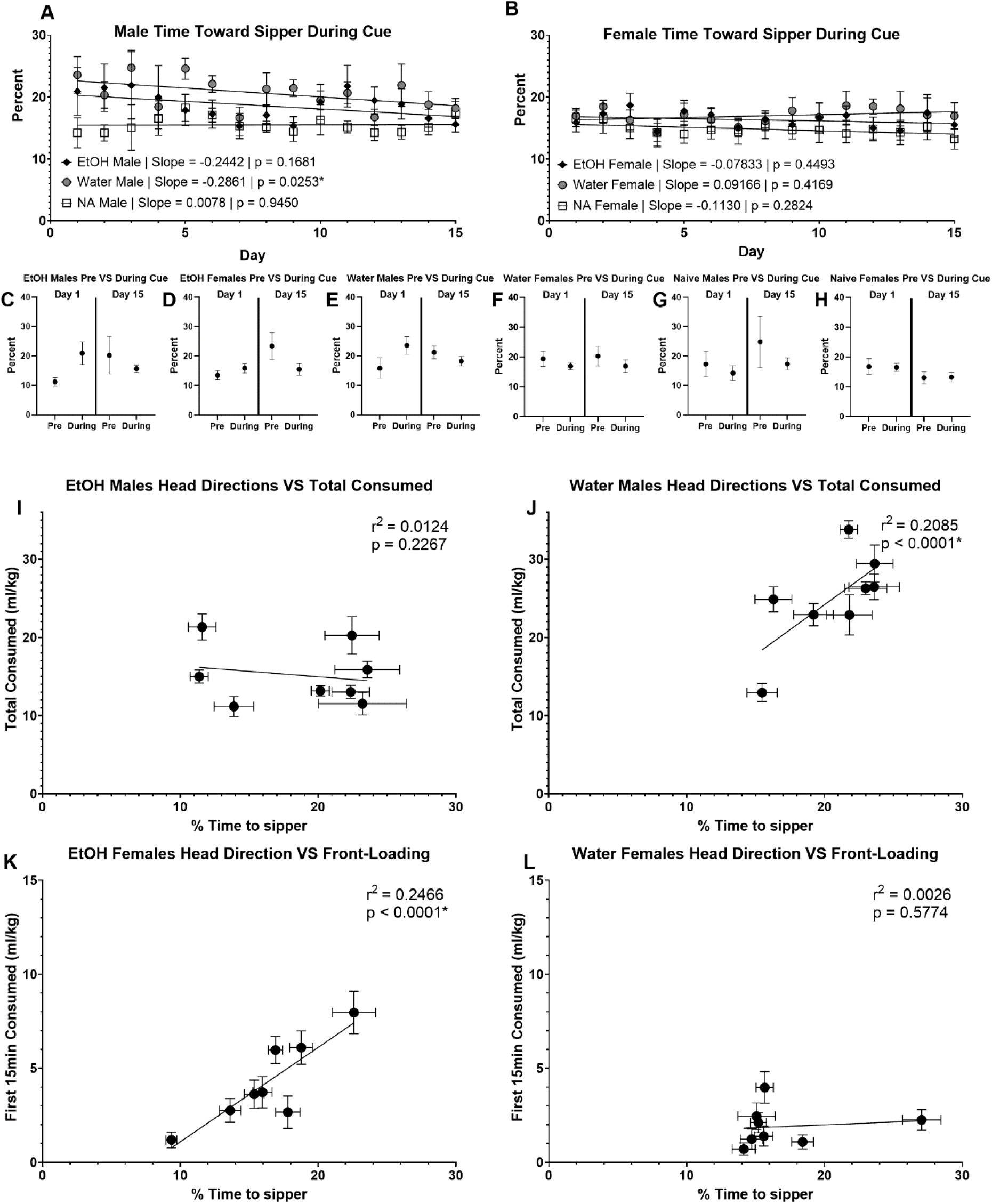
Impact of cues on head direction and relationship to fluid consumption. (A-B) The percent of time during the 15-minute cue period spent oriented toward the sipper, with “oriented” defined as a 90-degree angle extending out directly from the nose of the mouse. (C-H) t-tests of the percent of time spent oriented toward the sipper during the 15-minutes prior to the cue and during the 15-minute cue on day 1 and day 15, split by treatment and sex. (C) EtOH Males Day 1: df = 7, t = 2.259, p = 0.0584; Day 15: df = 7, t = 0.6929, p = 0.5107. (D) EtOH Females Day 1: df = 7, t = 0.9999, p = 0.3507; Day 15: df = 7, t = 2.083, p = 0.0758. (E) Water Males Day 1: df = 7, t = 1.392, p = 0.2066; Day 15: df = 7, t = 1.111, p = 0.3031. (F) Water Females Day 1: df = 7, t = 1.118, p = 0.3003; Day 15: df = 7, t = 0.8925, p = 0.4018. (G) Naïve Males Day 1: df = 7, t = 0.6607, p = 0.5299; Day 15: df = 7, t = 1.059, p = 0.3247. (H) Naïve Females Day 1: df = 7, t = 0.0725, p = 0.9442; Day 15: df = 7, t = 0.09, p = 0.9308. (I-L) Double error bar correlations between the percent of time oriented toward the sipper during the cue, total volume consumed during DID, and volume consumed during the first 15 minutes of DID.

No group experienced a significant change in the amount of time spent oriented toward the sipper during the cue as compared to the 15-minutes immediately prior to the cue on either day 1 or day 15 (Fig 3.C-H; p > 0.05).

Time spent oriented toward the sipper during the 15-minute cue predicted drinking patterns in a sex specific manner. In alcohol consuming females, this behavior predicted front-loading (Fig. 3.K; r2 = 0.2466, p < 0.0001), which was not observed in water consuming females (Fig. 3.L; r2 = 0.0026, p < 0.5574). Additionally, time spent oriented toward the sipper predicted the total volume consumed during 2-hours of DID in water males (Fig. 3.J; r2 = 0.2085, p < 0.0001), but not alcohol consuming males (Fig. 3.I; r2 = 0.0124, p = 0.2267). The relationships between head direction and consumption in other comparisons made were either statistically insignificant or very weak (Sup 1. A-D).

To further understand this relationship, we looked at the correlation between head direction and front-loading in each individual alcohol-consuming female mouse as a function of day (Sup. I-L). The relationship between frontloading and head direction varied greatly within a given subject, with only one mouse demonstrating a statistically significant correlation in the expected positive direction. However, there was great variability between subjects in average head direction, which was responsible for driving the significant correlation between frontloading and head direction between subjects reported above. This suggests that innate difference in head direction during cue presentation predicted the development of frontloading.

### 3.4 Distance from the sipper

The last cue period behavior analyzed was the distance from the sipper site. A 3-way ANOVA revealed main effects of sex (DF = 1, F = 6.941, p = 0.0117), treatment (DF = 2, F = 3.482, p = 0.0399), and day (DF = 14, F = 14.151, p < 0.0001), in addition to a three-way interaction (DF = 28, F = 1.674, P = 0.0171). Regression analysis further demonstrated the complicated nature of this relationship, with all groups displaying significant decreases in the distance from the sipper during the cue except alcohol and Naïve males (Fig. 4A-B).

**Figure 4.**
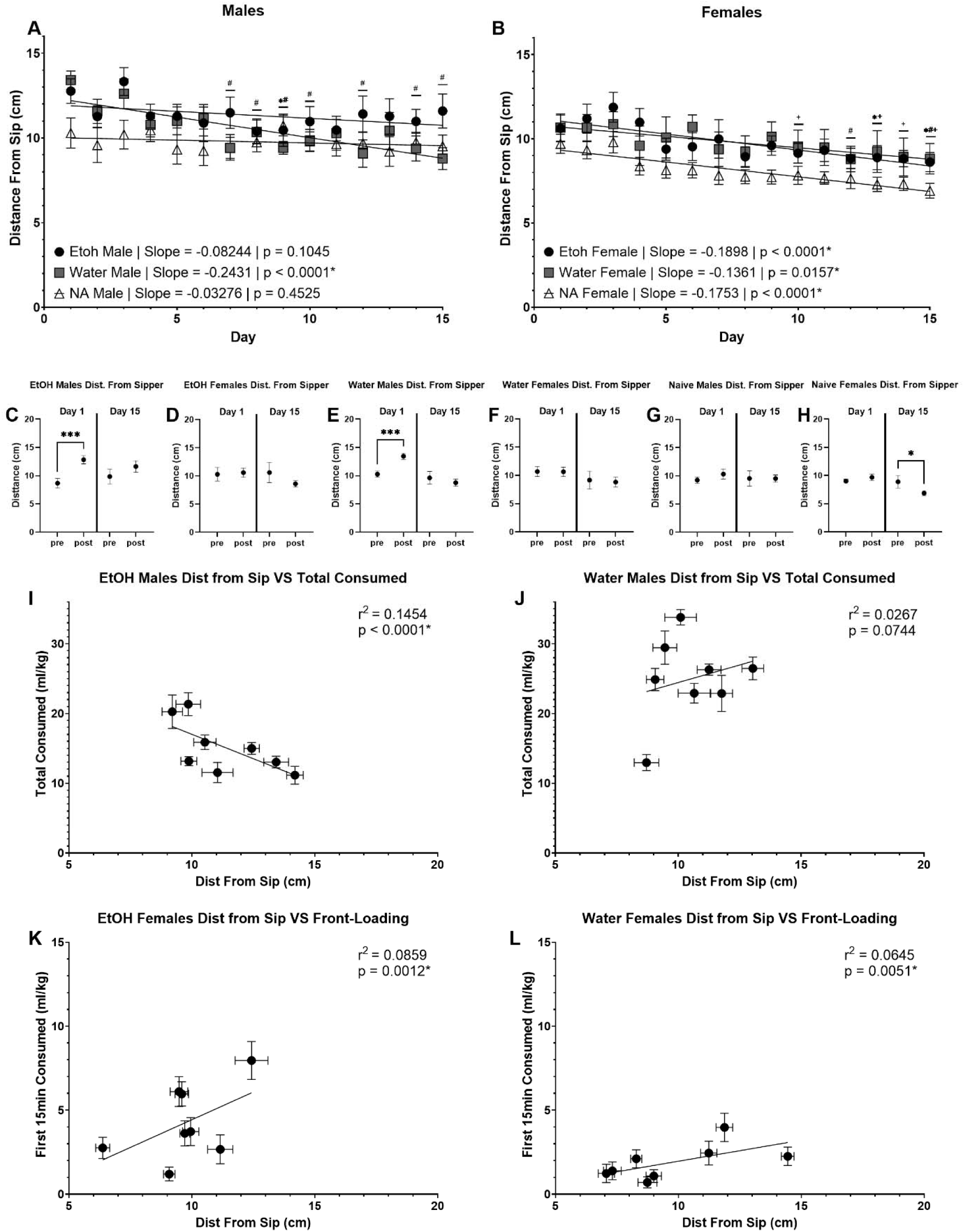
Impact of cues on distance from sipper and relationship to fluid consumption. (A-B) Average distance from the sipper during the 15-minute cue period, split by treatment and sex. “*” represents ethanol, “#” represents water, and “+” represents Naïve mice in a 2-way ANOVA Dunnett’s multiple comparisons test comparing each day to day 1. (C-H) t-tests of average distance from the sipper 15-minutes prior to the cue and during the 15-minute cue on day 1 and day 15, split by treatment and sex. (C) EtOH Males Day 1: df = 7, t = 7.764, p = 0.0001; Day 15: df = 7, t = 1.509, p = 0.175. (D) EtOH Females Day 1: df = 7, t = 0.4715, p = 0.6516; Day 15: df = 7, t = 1.476, p = 0.1834. (E) Water Males Day 1: df = 7, t = 5.747, p = 0.0007; Day 15: df = 7, t = 0.7891, p = 0.4559. (F) Water Females Day 1: df = 7, t = 0.0488, p = 0.9625; Day 15: df = 7, t = 0.3423, p = 0.7422. (G) Naïve Males Day 1: df = 7, t = 1.933, p = 0.0944; Day 15: df = 7, t = 0.0083, p = 0.9936. (H) Naïve Females Day 1: df = 7, t = 0.9941, p = 0.3533; Day 15: df = 7, t = 2.417, p = 0.0463. (I-L) Double error bar correlations between the average distance from the sipper during the cue, total volume consumed during DID, and volume consumed during the first 15 minutes of DID.

On day 1, alcohol and water consuming males reacted to the cue by moving significantly further away from the sipper as compared to the 15-minutes immediately prior to its presentation (Fig 4.C, E; p < 0.05). On day 15, Naïve females moved significantly closer to the sipper during the cue as compared to prior to its presentation (Fig 4.H; p < 0.05).

Distance from the sipper was also predictive of consumption patterns in a sex specific manner. The strongest relationship, albeit of moderate effect size, was with total volume consumed in alcohol males (Fig. 4.I; r2 = 0.1454, p < 0.0001) which was not observed in water males (Fig. 4.J; r2 = 0.0267, p =0.0744). Additionally, there were weak but statistically significant positive relationships between distance from the sipper and front-loading in females of both treatments (Fig. 4.K-L; alcohol: r2 = 0.0859, p = 0.0012; water: r2= 0.0646, p = 0.0051). The relationships between distance from the sipper during the cue and total consumption in females and front-loading in males were either statistically insignificant or weak (Sup.1 E-H).

## 4. Discussion

This work represents a uniquely detailed behavioral analysis within a Pavlovian conditioning paradigm involving home-cage testing, temporally precise fluid intake measurements, high-resolution behavioral tracking, and remotely presented cues. We hypothesized that with daily cue-sipper pairings, only mice receiving alcohol would orient toward and approach the sipper port during cues predicting its availability. Contrary to expectations, the only consistent cue-related change across sessions was an increase in locomotor activity, observed in all groups, including those that never received specialized sippers. Nonetheless, certain cue-period behaviors were correlated with specific alcohol consumption elements. Among alcohol-consuming females, those with higher time oriented toward the sipper port during the auditory cue displayed greater front-loading of alcohol. In males, closer proximity to the sipper during the cue period correlated with higher total alcohol intake. Although these associations were modest in strength, they suggest potential sex-specific behavioral predictors of alcohol consumption. Indeed, prior work has indicated that various other measures of cue-reactivity and motivation, such as sign-tracking, Pavlovian approach, and Pavlovian-Instrumental Transfer, may sex-dependently predict consumption in rodents (Barker & Taylor, 2019).

To this end, previous investigations into cue-alcohol associative learning in rodents have primarily relied on operant or similar behavioral testing chambers. While well-suited for detecting goal-or sign-tracking behaviors, particularly when visual cues are spatially distinct from the location of reward receipt (Krank, 2003; Krank et al., 2008), these environments necessarily introduce new contextual cues and handling-induced stress that can potentially impact behavioral interpretations (Balcombe et al., 2004). Furthermore, the experimenter must first go into the vivarium where subjects are housed, relocate them, and then interact with them physically to deposit into testing chambers. Thus, the experimenter provides an array of salient cues that predict the availability of reward far in advance of chamber cues. We therefore argue here that the experimenter may be the most salient cue in these conditions, and that (non-recorded) behavioral responses during these interactions may occur that are more correlated to the current motivational state of the subject than those recorded during operant testing.

Nonetheless, our results are in line with one study conducted in operant chambers that found that cue-induced approach to the sipper predicted alcohol consumption and lick rate (Cofresí et al., 2018), but not another where no relationship was observed between operant responses for alcohol during presentation of discrete cues and blood alcohol concentration (Blegen et al., 2018).

While our main findings support the view that cue-induced behaviors predict the motivational state of the animal, it is important to consider caveats to this interpretation. First, we did not observe evidence of cue-induced learning-related behavioral changes. We also did not observe changes in head direction in response to the cues compared to baseline. Thus, the relationship we observed in females between head direction during cues and frontloading supports pre-existing individual differences in cue-reactivity that predisposes these subjects to drinking alcohol more quickly. Alternatively, head direction during cues may co-occur with faster alcohol consumption due to an unidentified third variable influencing both. Clearly further research is warranted to disentangle innate from learned responses in the context of our experimental design.

While remote cue initiation guaranteed auditory cues were experienced prior to those associated with human experimenters, human interaction was not entirely eliminated. Fully automated designs devoid of human interaction would further minimize confounding variables. Additionally, the auditory cue in our design was considerably longer than is typical in delay conditioning paradigms, which may have resulted in more robust conditioning. Furthermore, the ‘clicking’ noise serving as the cue is produced by the VDM system when a bead of fluid is dispensed. Therefore, the same noise serving as the cue was active intermittently throughout the 2-hour drinking period contingent on mouse consumption. This design reduced the temporal specificity of conditioned responses and potentially blurred the distinction between discrete and contextual cues. Indeed, it has been found that with repeated cue-alcohol pairings within a single 35-minute time window, the level of cue-induced approach to the sipper location decays in male (Cofresí, Grote, et al., 2019), but not female rats (Cofresí, Monfils, et al., 2019). Among females, this same group found a moderate correlation between cue-induced sipper location approach and BEC. Given this, it is possible that our cue period was long enough to produce a decay in response within single cue-sipper presentations, potentially dampening our effects and driving sex differences. Their team was also able to show evidence of associative learning with a visual delay cue predicting alcohol availability, but this cue was not necessary for the development of alcohol seeking. Instead, they posit that the introduction of the sipper itself may be a more salient cue driving rapid consummatory behavior. Similarly, this may be the case in the present study, with the introduction of the sipper being a more salient cue than the ongoing auditory stimuli. Additionally, earlier work found that CS-US interval is important for the development of conditioned magazine approach for appetitive rewards (sucrose and pellets), with shorter intervals promoting higher levels of approach (Delamater & Holland, 2008).

Therefore, measuring reactions to a long complex cue as opposed to the recurring stimuli immediately prior to sipper receipt may obscure our ability to identify behavioral markers of risky drinking behavior. Therefore, shortening the cue period in future studies could enable more precise analyses of rapid, pose-specific behavioral changes and allow for closer comparison with the existing classical conditioning literature.

Currently, we are integrating this behavioral framework with *in vivo* neuroscience techniques. By temporally aligning behavioral output with neural data captured via electrophysiology, fiber photometry, etc., we are positioned to conduct closed-loop experimental designs, in which optogenetic manipulations are delivered in real-time based on predictive behavioral signatures to reduce alcohol consumption. Such approaches could yield powerful insights into the neural mechanisms of motivation for alcohol and offer a path toward individualized interventions for excessive alcohol use.

In conclusion, this study serves as a first step in demonstrating that temporally precise behavioral tracking in a low-intervention home-cage setting, can reveal distinct, cue-elicited responses predictive of excessive alcohol consumption. By limiting contextual confounds, utilizing remote cue initiation, and leveraging scalable, automated measurements, this model offers a valuable tool for identifying early behavioral signatures of binge-like drinking, and can relatively easily be paired with mechanistic studies using in-vivo systems neuroscience techniques.

## Author Contributions

SAD, DAF, MO, and DNL performed data collection. Analysis was led by SAD and DNL. Conceptualization was performed by SAD, DAF, MO, and DNL.

## Acknowledgments

Nicole M. Maphis^1^-Proof-reading/editing

