## Supplementary figures and images for "Remotely Presenting Alcohol-predicting Cues Avoids Confound of Experimenter as First Cue and Reveals Sex-specific Behaviors that Predict the Rate and Amount of Alcohol Consumption"

### Supplementary_Figures

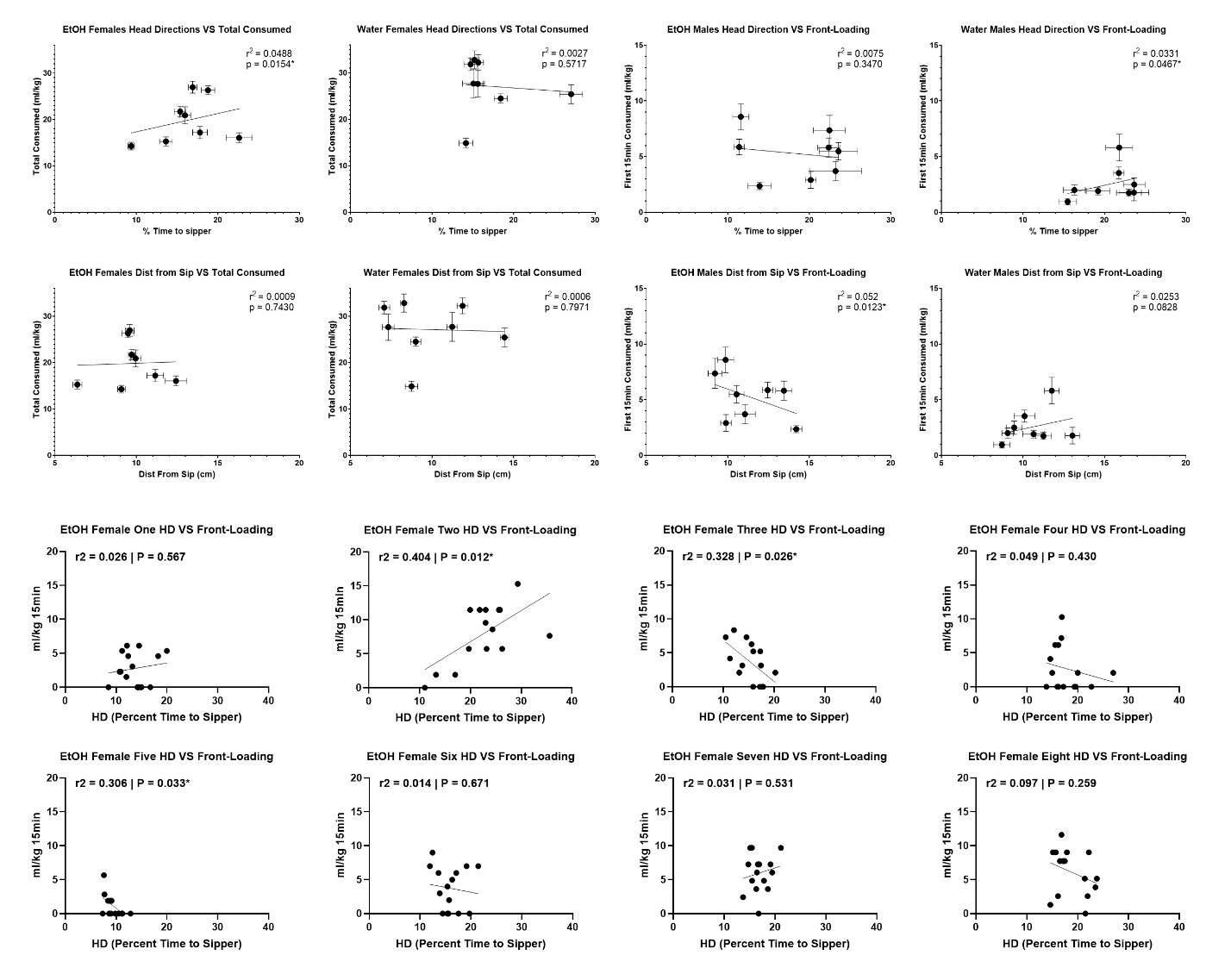

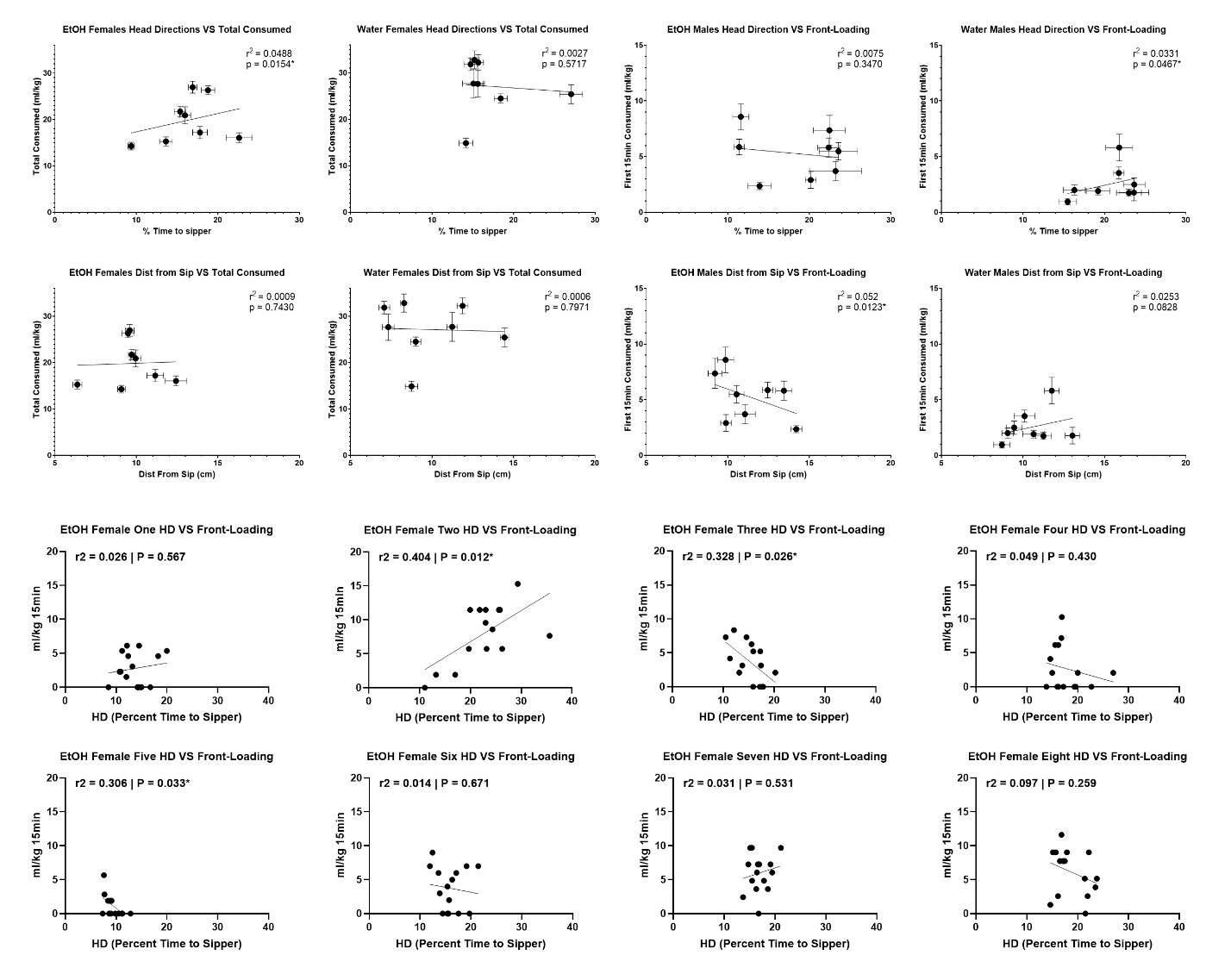
